# An *in vivo* binding assay for RNA-binding proteins based on repression of a reporter gene

**DOI:** 10.1101/168625

**Authors:** Noa Katz, Roni Cohen, Oz Solomon, Beate Kaufmann, Noa Eden, Orna Atar, Zohar Yakhini, Sarah Goldberg, Roee Amit

## Abstract

We employ a reporter assay and Selective 2′-hydroxyl acylation analysed by primer extension sequencing (SHAPE-seq) to study translational regulation by RNA-binding proteins, in bacteria. We designed 82 constructs, each with a single hairpin based on the binding sites of the RNA-binding coat proteins of phages MS2, PP7, GA, and Qβ, at various positions within the N-terminus of a reporter gene. In the absence of RNA-binding proteins, the translation level depends on hairpin location, and exhibits a three-nucleotide periodicity. For hairpin positions within the initiation region, we observe strong translational repression in the presence of its cognate RNA-binding protein. *In vivo SHAPE-seq* results for a representative construct indicate that the repression phenomenon correlates with a wide-swath of protection, including the hairpin and extending past the ribosome binding site. Consequently, our data suggest that the protection provided by the RBP-hairpin complex inhibits ribosomal initiation. Finally, utilizing the repression phenomenon for quantifying protein-RNA binding affinity *in vivo*, we both observe partially contrasting results to previous *in vitro* and *in situ* studies, and additionally, show that this method can be used in a high-throughput assay for a quantitative study of protein-RNA binding *in vivo*.

## INTRODUCTION

The regulation of gene expression is a process central to all biological life-forms. It is a process thought to be mediated largely by proteins, which interact with either chromatin or its RNA product. The best-known form of regulation is mediated by transcription factors, which control RNA levels by their sequence-specific interaction with DNA. Post transcriptional regulation based on protein-RNA interactions, however, is quite different, due to the nature of RNA. Unlike DNA which is a long, chromatinized, replicated, and for the most part exists as a double stranded molecule, RNA is a short, transient (i.e. constantly manufactured and degraded), exists in multiple copies, has particular modifications (1, 2), and folds into functional secondary and tertiary structures. RNA structure is thought to be highly dynamic, and is dependent on many factors such as temperature, cellular RNA-binding protein (RBP) content, presence or absence of translating ribosomes, and interaction with other RNA molecules (3). Thus, a typical RBP-RNA interaction is likely to depend not only on the presence of a specific binding sequence, but also on many other factors.

In bacteria, post-transcriptional regulation has been studied extensively in recent decades. There are well-documented examples of RBPs that either inhibit or directly compete with ribosome binding via a variety of mechanisms. These include direct competition with the 30S ribosomal subunit for binding via single stranded recognition (4), entrapment of the 30S subunit in an inactive complex via a nested pseudoknot structure (5) and ribosome assembly inhibition when the RBP is bound to a structured RBP binding site, or hairpin (6–9). These hairpins have been studied in three distinct positions: either immediately downstream and including the AUG (7), upstream of the Shine-Dalgarno sequence (8), or as structures that entrap Shine-Dalgarno motifs, as is the case for the PP7 and MS2 phage coat-protein binding sites. There is also a well-characterized example of translation stimulation: binding of the phage Com RBP was shown to destabilize a sequestered ribosome binding site (RBS) of the Mu phage *mom* gene, thereby facilitating translation (10, 11). While these studies indicate a richness of RBP-RNA-based regulatory mechanisms, a systematic understanding of the relationship between RBP binding, sequence specificity, the underlying secondary and tertiary RNA structure, and the resultant regulatory output is still lacking.

In recent years, advances in next generation sequencing (NGS) technology combined with selective nucleic acid labeling approaches have facilitated focused study of specific RNA structures *in vivo*. These chemical-modification approaches (12–16) can generate a “foot-print” of the dynamical structure of a chosen mRNA molecule *in vivo*, while in complex with ribosomes and/or other RBPs. In parallel, synthetic biology approaches that simultaneously characterize large libraries of synthetic regulatory constructs have been increasingly used to complement the detailed study of single mRNA transcripts. While these synthetic approaches have been mostly applied to the transcriptional regulatory platforms (17–20), their potential for deciphering post-transcriptional regulatory mechanisms have been demonstrated in a recent study that interrogated IRES sequences in mammalian cells (21).

Building on these advances and on a smaller-scale demonstration of translational repression by the RBP L7Ae in both bacteria and mammalian cells (22), we measured the regulatory output of a small library of synthetic constructs in which we systematically varied the position and type of RBP binding sites. In addition, we applied Selective 2′-hydroxyl acylation analysed by primer extension sequencing (SHAPE-seq) (23, 24, 15) to a single variant, to further identify and characterize RBP-based regulatory mechanisms in bacteria. Our findings indicate that the chosen structure-binding RBPs (coat proteins from the bacteriophages GA (25), MS2 (26), PP7 (9), and Qβ (27)), generate a strong repression response when bound to the translation initiation region. This inhibitory response is associated with a strong protection effect that spans a large segment of the RNA that includes both the binding site and the RBS. We employed this strong repression phenomenon as an *in vivo* binding assay for RBP-RNA interactions. Using this assay, we quantitatively characterized RBP binding to a set of mutated binding sites in a high-throughput manner, thereby increasing our understanding of RBP-RNA binding *in vivo* and enabling the engineering of more complex RNA-based applications.

## MATERIALS AND METHODS

### Design and construction of binding-site plasmids

Binding-site cassettes (see Supplementary Table 1) were ordered either as double-stranded DNA minigenes from Gen9 or as cloned plasmids (minigene + vector) from Twist Biosciences. Each minigene was ∼500 bp long and contained the parts in the following order: Eagl restriction site, ∼40 bases of the 5’ end of the Kanamycin (Kan) resistance gene, pLac-Ara promoter, ribosome binding site (RBS), an RBP binding site, 80 bases of the 5’ end of the mCherry gene, and an ApaLI restriction site. As mentioned, each cassette contained either a wild-type or a mutated RBP binding site (see Supplementary Table 1), at varying distances downstream to the RBS. All binding sites were derived from the wild-type binding sites of the coat proteins of one of the four bacteriophages MS2, PP7, GA and Qβ. For insertion into the binding-site plasmid backbone, they were double-digested with Eagl-HF and ApaLI (New England Biolabs [NEB]). The digested minigenes were then cloned into the binding-site backbone containing the rest of the mCherry gene, terminator, and a Kanamycin resistance gene, by ligation and transformation into *E. coli* TOP10 cells (ThermoFisher Scientific). Purified plasmids were stored in 96-well format, for transformation into *E. coli* TOP10 cells containing one of four fusion-RBP plasmids (see below).

### Design and construction of fusion-RBP plasmids

RBP sequences lacking a stop codon were amplified via PCR off of either Addgene or custom-ordered templates (Genescript or IDT, see Supplementary Table 2). All RBPs presented (MCP, PCP, GCP, and QCP) were cloned into the RBP plasmid between restriction sites KpnI and AgeI, immediately upstream of an mCerulean gene lacking a start codon, under the pRhlR promoter (containing the *rhlAB* las box (28)) and induced by C_4_-HSL. The backbone contained an Ampicillin (Amp) resistance gene. The resulting fusion-RBP plasmids were transformed into *E. coli* TOP10 cells. After Sanger sequencing, positive transformants were made chemically-competent and stored at −80°C in 96-well format.

### Transformation of binding-site plasmids

Binding-site plasmids stored in a 96-well format were simultaneously transformed into chemically-competent bacterial cells containing one of the RBP-mCeulean plasmids. After transformation, cells were plated using an 8-channel pipettor on 8-lane plates (Axygen) containing LB-agar with relevant antibiotics (Kan and Amp). Double transformants were selected, grown overnight, and stored as glycerol stocks at −80°C in 96-well plates (Axygen).

### RNA extraction and reverse-transcription for qPCR measurements

Starters of *E. coli* TOP10 containing the relevant constructs on plasmids were grown in LB medium with appropriate antibiotics overnight (16 hr). The next morning, the cultures were diluted 1:100 into fresh semi-poor medium and grown for five hours. For each isolation, RNA was extracted from 1.8 ml of cell culture using standard protocols. Briefly, cells were lysed using Max Bacterial Enhancement Reagent followed by TRIzol treatment (both from Life Technologies). Phase separation was performed using chloroform. RNA was precipitated from the aqueous phase using isopropanol and ethanol washes, and then resuspended in RNase-free water. RNA quality was assessed by running 500 ng on 1% agarose gel. After extraction, RNA was subjected to DNAse (Ambion/Life Technologies) and then reverse-transcribed using MultiScribe Reverse Transcriptase and random primer mix (Applied Biosystems/Life Technologies). For qPCR experiments, RNA was isolated from three individual colonies for each construct.

### qPCR measurements

Primer pairs for mCherry and normalizing gene *idnT* were chosen using the Primer Express software and aligned using BLAST (29) (NCBI) with respect to the *E. coli* K-12 substr. DH10B (taxid:316385) genome (which is similar to TOP10) to avoid off-target amplicons. qPCR was carried out on a QuantStudio 12K Flex machine (Applied Biosystems/Life Technologies) using SYBR-Green. Three technical replicates were measured for each of the three biological replicates. A *C_T_* threshold of 0.2 was chosen for all genes.

### *in vivo* SHAPE-seq

LB medium supplemented with appropriate concentrations of Amp and Kan was inoculated with glycerol stocks of bacterial strains harboring both the binding-site plasmid and the RBP-fusion plasmid (see Supplementary Table 3 for details of primers and barcodes, and Supplementary Figure 2), and grown at 37°C for 16 hours while shaking at 250 rpm. Overnight cultures were diluted 1:100 into semi-poor medium. Each bacterial sample was divided into a non-induced sample and an induced sample in which RBP protein expression was induced with 250 nM N-butanoyl-L-homoserine lactone (C_4_-HSL), as described above.

Bacterial cells were grown until OD_600_=0.3, 2 ml of cells were centrifuged and gently resuspended in 0.5 ml semi-poor medium supplemented with a final concentration of 30 mM 2-methylnicotinic acid imidazole (NAI) suspended in anhydrous dimethyl sulfoxide (DMSO, Sigma Aldrich) (15, 23), or 5% (v/v) DMSO. Cells were incubated for 5 min at 37°C while shaking and subsequently centrifuged at 6000 g for 5 min. Column-based RNA isolation (RNeasy mini kit, QIAGEN) was performed for the strain harboring PP7-wt δ=6. Samples were divided into the following sub-samples (Supplementary Figure 2A):

1. induced/modified (+C_4_-HSL/+NAI)
2. non-induced/modified (-C_4_-HSL/+NAI)
3. induced/non-modified (+C_4_-HSL/+DMSO)
4. non-induced/non-modified (-C_4_-HSL/+DMSO).

Subsequent steps of the SHAPE-seq protocol, that were applied to all samples, have been described elsewhere (24), including reverse transcription (steps 40-51), adapter ligation and purification (steps 52-57) as well as dsDNA sequencing library preparation (steps 68-76). In brief, 1000 ng of RNA were converted to cDNA using the reverse transcription primers (for details of primer and adapter sequences used in this work see Supplementary Table 3). The RNA was mixed with 0.5 µM primer for mCherry (#1) and incubated at 95°C for 2 min followed by an incubation at 65°C for 5 min. The Superscript III reaction mix (Thermo Fisher Scientific; 1x SSIII First Strand Buffer, 5 mM DTT, 0.5 mM dNTPs, 200 U Superscript III reverse transcriptase) was added to the cDNA/primer mix, cooled down to 45°C and subsequently incubated at 52°C for 25 min. Following inactivation of the reverse transcriptase for 5 min at 65°C, the RNA was hydrolyzed (0.5 M NaOH, 95°C, 5 min) and neutralized (0.2 M HCl). cDNA was precipitated with 3 volumes of ice-cold 100% ethanol, incubated at −80°C for 15 minutes, centrifuged at 4°C for 15 min at 17000 g and resuspended in 22.5 µl ultra-pure water. Next, 1.7 µM of 5’ phosphorylated ssDNA adapter (#2) (see Supplementary Table 3) was ligated to the cDNA using a CircLigase (Epicentre) reaction mix (1xCircLigase reaction buffer, 2.5 mM MnCl_2_, 50 µM ATP, 100 U CircLigase). Samples were incubated at 60°C for 120 min, followed by an inactivation step at 80°C for 10 min. cDNA was ethanol precipitated (3 volumes ice-cold 100% ethanol, 75 mM sodium acetate [pH 5.5], 0.05 mg/mL glycogen [Invitrogen]). After an overnight incubation at −80°C, the cDNA was centrifuged (4°C, 30 min at 17000 g) and resuspended in 20 µl ultra-pure water. To remove non-ligated adapter (#2), resuspended cDNA was further purified using the Agencourt AMPure XP beads (Beackman Coulter) by mixing 1.8x of AMPure bead slurry with the cDNA and incubation at room temperature for 5 min. The subsequent steps were carried out with a DynaMag-96 Side Magnet (Thermo Fisher Scientific) according to the manufacturer’s protocol. Following the washing steps with 70% ethanol, cDNA was resuspended in 20 μl ultra-pure water. cDNAs were subjected to PCR amplification to construct dsDNA library as detailed below.

### RBP protection assay using *in vitro* SHAPE-seq

*In vitro* modification was carried out on non-induced, DMSO-treated samples (Supplementary Figure 2A) and has been described elsewhere (15). Briefly, 1500 ng of isolated RNA were denatured at 95°C for 5 min, transferred to ice for 1 min and incubated in SHAPE-seq reaction buffer (100 mM HEPES [pH 7.5], 20 mM MgCl2, 6.6 mM NaCl) supplemented with 40 U of RiboLock RNase inhibitor (Thermo Fisher Scientific) for 5 min at 37°C allowing the RNA molecule to refold. Next, we added 15.6 pmol (based on 1:2 molar ratio between RNA:PP7 protein) of highly-purified recombinant PP7 protein (Genscript) to the RNA samples and incubated at 37°C for 30 min. Subsequently, final concentrations of 100 mM NAI or 5% (v/v) DMSO were added to the RNA-PP7 protein reaction mix and incubated for an additional 10 min at 37°C. Samples were then transferred to ice to stop the SHAPE reaction and precipitated by addition of 300 µl ice-cold 100% ethanol, 10 µl Sodium Acetate 3M, 0.5 µl ultrapure glycogen (Thermo scientific) and 70 µl DEPC-treated water. Samples were incubated at −80°C for 15 min followed by centrifugation at 4°C, 17000 g for 15 min. Supernatant was removed and samples were air-dried for 5 min at room temperature and resuspended in 10 μl of RNAse-free water.

### SHAPE-Seq library preparation and sequencing

To produce the dsDNA for sequencing 10ul of purified cDNA from the SHAPE procedure (see above) were PCR amplified using 3 primers: 4nM mCherry selection (#3) (primer extends 4 nucleotides into mCherry transcript to avoid the enrichment of ssDNA-adapter products), 0.5µM TruSeq Universal Adapter (#4) and 0.5µM TrueSeq Illumina indexes (one of #5-16) (Supplementary Table 3) with PCR reaction mix (1x Q5 HotStart reaction buffer, 0.1 mM dNTPs, 1 U Q5 HotStart Polymerase [NEB]). A 15-cycle PCR program was used: initial denaturation at 98°C for 30 s followed by a denaturation step at 98°C for 15 s, primer annealing at 65°C for 30 s and extension at 72°C for 30 s, followed by a final extension 72°C for 5 min. Samples were chilled at 4°C for 5 min. After cool-down, 5 U of Exonuclease I (ExoI, NEB) were added, incubated at 37°C for 30 min followed by mixing 1.8x volume of Agencourt AMPure XP beads to the PCR/ExoI mix and purified according to manufacturer’s protocol. Samples were eluted in 20 µl ultra-pure water. After library preparation, samples were analyzed using the TapeStation 2200 DNA ScreenTape assay (Agilent) and the molarity of each library was determined by the average size of the peak maxima and the concentrations obtained from the Qubit fluorimeter (Thermo Fisher Scientific). Libraries were multiplexed by mixing the same molar concentration (2-5 nM) of each sample library and sequenced using the Illumina HiSeq 2500 sequencing system using 2x100 bp paired-end reads.

### Analysis routines and models

See Supplementary Information.

## RESULTS

### Hairpin inhibition of translation varies as a function of position inside coding region

We first studied the translational regulatory effects of hairpins located within the 5’ end of a transcribed gene. We chose to work with RBP binding-site sequences encoding hairpin structure (Fig. 1A, all structures herein were generated using NUPACK (30)), in the absence of their cognate RBPs [the phage coat proteins of bacteriophages MS2 (MCP), PP7 (PCP), GA (GCP), and Qβ (QCP)]. The four wild-type binding sites are characterized by two stems of varying length, which are separated by a single unpaired nucleotide or "bulge”, and a loop of differing size that varies from 3 to 6 nucleotides (Fig. 1A). We measured the effect of hairpin position using a reporter system comprising a plasmid with a fluorescent mCherry gene under a constitutive pLac/Ara promoter (Fig. 1B-top), with a single binding site inserted into the mCherry coding region (additional nucleotides were inserted downstream of the binding site to maintain the reading frame). Hairpin position δ measured from the first nucleotide of the mCherry AUG was varied from 4 (immediately adjacent to the AUG) to 12 nt, at 1 nt resolution excluding positions that put a stop codon or an AUG in frame, and from 5 to 21 nt for PP7-wt and GA-wt (see Supplementary Table 1 for sequences). In Fig. 1B, we plot the mean expression level of each construct as a function of d. The data show that mCherry expression depends strongly on the position of the hairpin relative to the AUG, varying between inhibition to strong expression, relative to the expression level for unstructured mRNA (black dashed line, averaged over two variants lacking apparent hairpins at the N-terminus; see Supplementary Table 1 – one with only the natural mCherry reporter, and another with a codon for Glycine [GGC] added after the AUG). In particular, constructs containing the Qβ-wt and GA-wt hairpins exhibit approximately a two-order-of-magnitude increase in expression when δ is shifted from 5 to 6, and 4 to 5, respectively.

**Fig. 1:**
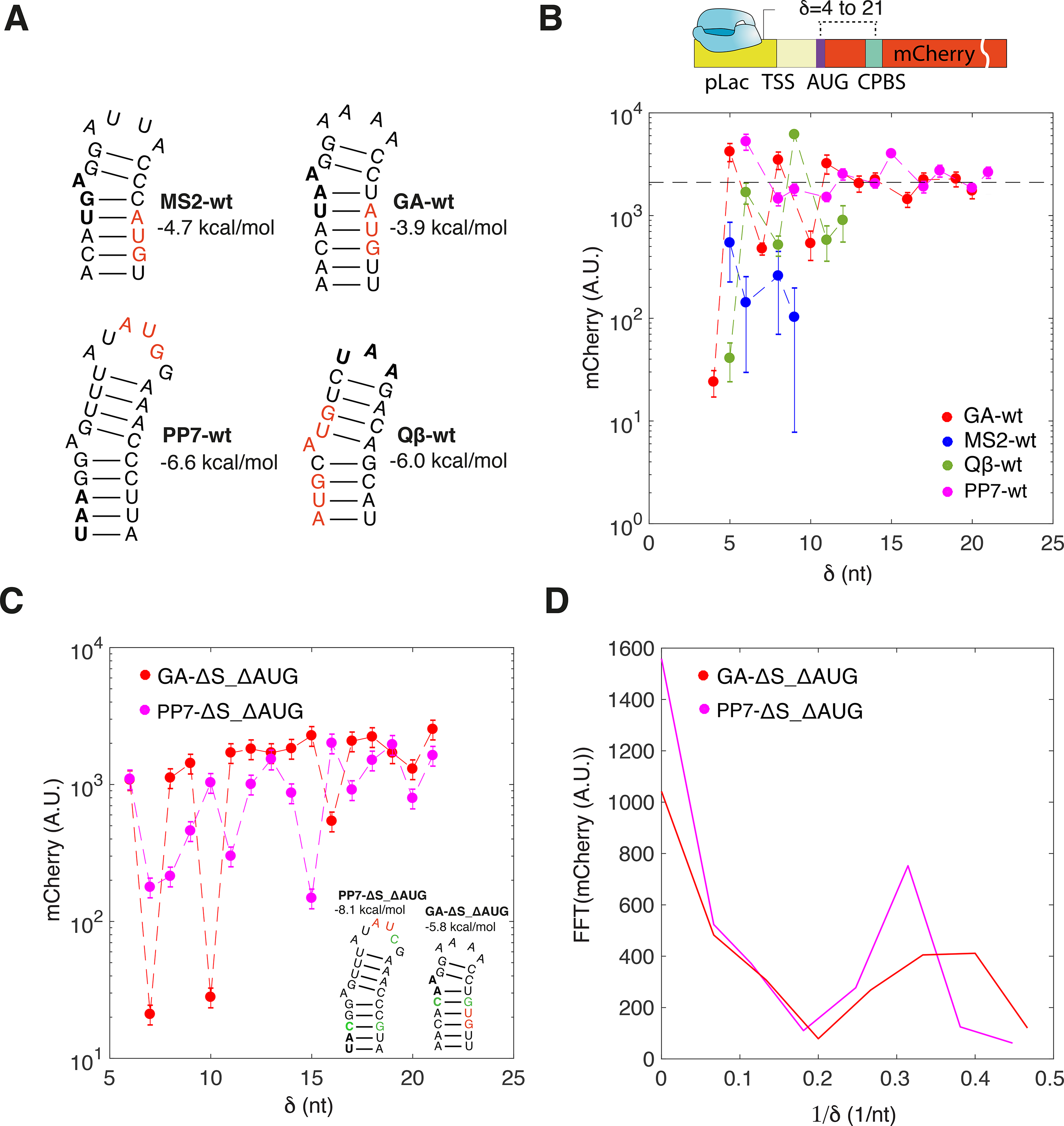
Dependence of mCherry expression level on hairpin position. (A) The four hairpins used in this experiment were the native (wt) binding sites for the MS2, PP7, GA, and Qβ coat proteins. Stop codons and start codons inside the binding sites are highlighted, in bold and red. (B) Mean mCherry basal expression levels for 26 variants as a function of distance from the A of the AUG. Data for PP7-wt (magenta), GA-wt (red), Qβ-wt (green), and MS2-wt (blue) binding sites as function of hairpin position d, in the absence of RBPs (non-induced case). Data shown only for positions where the internal AUG and stop codons are out of-frame, for PP7, MS2, GA and the first AUG only, for Qβ; mCherry levels for all proteins for the in-frame positions were found to be effectively zero (data not shown). For Qβ-wt positions 6,9, and 12 the second internal AUG may also be active in initiation contributing to a higher translational levels. Dashed line represents the average basal level for two constructs with a non-structured sequence replacing the binding site downstream of the AUG. CPBS - stands for coat-protein binding site (schemas, top). (C) Mean mCherry basal production rate for 34 variants as a function of distance (d) from the A of the AUG. Data for PP7-ΔS_ΔAUG (magenta) and GA-ΔS_ΔAUG (red). (Inset) structures of the hairpins with mutations colored in green. (D) FFT analysis on the rate of production data displayed in (C) with the magneta and red FFT traces corresponding to the position-based rate of production data for the PP7-ΔS_ΔAUG and GA-ΔS_ΔAUG constructs respectively.

Interestingly, mCherry expression for all four hairpins seems to suggest a 3-nt periodicity, indicating that the basal expression levels may depend on the position of the hairpin within the reading frame of the translating ribosome. However, all four hairpins contain both a start and stop codons (Fig. 1A – red and bold letters respectively), which can affect the level of expression and, therefore, preclude us from exploring the positions where they are located in-frame with respect to the mCherry reporter gene. To determine whether this apparent 3-nt periodicity is a robust experimental feature of these hairpins, we constructed two mutated binding sites for PP7-wt and GA-wt, where the stop and start codons in each structure were eliminated (see Fig. 1C-inset). We then constructed 17 variants of each hairpin, altering their position within the reporter gene N-terminus at one nucleotide resolution starting from δ=6 to δ=21. For both mutated hairpins, the mean expression level data for each position exhibits an oscillatory behavior with an apparent 3-nt periodicity. To confirm this periodic signature, we carried out a fast Fourier transform analysis (FFT) on both expression level signals. The analysis shows that (Fig. 1D) a peak centered at frequency of 0.3-0.4 (1/nt) appears for both the PP7-ΔS_ΔAUG and GA-ΔS_ΔAUG constructs strengthening the 3-nt periodic observation. Finally, we note that for both constructs the amplitude of oscillations reduce sharply for positions where δ>15, indicating that this periodicity effect is likely to be related to initiation of translation.

### RBPs repress translation when bound within the ribosomal initiation region of mRNA

We next explored the regulatory effect of an RBP-hairpin complex, as a function of hairpin position downstream to the AUG. To do so, we constructed 22 two-plasmid strains (Fig. 2A): the first was a subset of the plasmids that were used for the hairpin experiments in the N-terminus region (see Fig. 1B-top for schematic and Supplementary Table 1), while the second encoded an RBP fused to mCerulean, under a pRhlR promoter inducible by N-butanoyl-L-homoserine lactone (C_4_-HSL). We used two different RBPs: the phage coat proteins QCP and PCP of bacteriophages Qβ and PP7, respectively (see Supplementary Table 2). Each RBP—binding-site plasmid pair was transformed into *E. coli* TOP10 and grown in 24 different C_4_-HSL concentrations, in duplicate. Optical density, mCherry, and mCerulean fluorescence levels were measured at multiple time points for each inducer concentration. From these data, mCherry production rates (31, 32) were computed over a 2-3 hours window (see Supplementary Methods and Supplementary Fig. 1) for each inducer level, and mCerulean levels were averaged over the same time-frames. In Fig. 2B, we plot a series of dose-response curves obtained for PCP on three constructs containing the PP7-wt binding site, positioned at δ−8 (red), 12 (blue), and 17 (green) nt. For the hairpin located at δ=8, the mCherry production rate is reduced by nearly two orders of magnitude, while the hairpin positioned at δ=12, produced a weakly repressing dose-response function, and no repression was observed at δ−17.

**Fig. 2:**
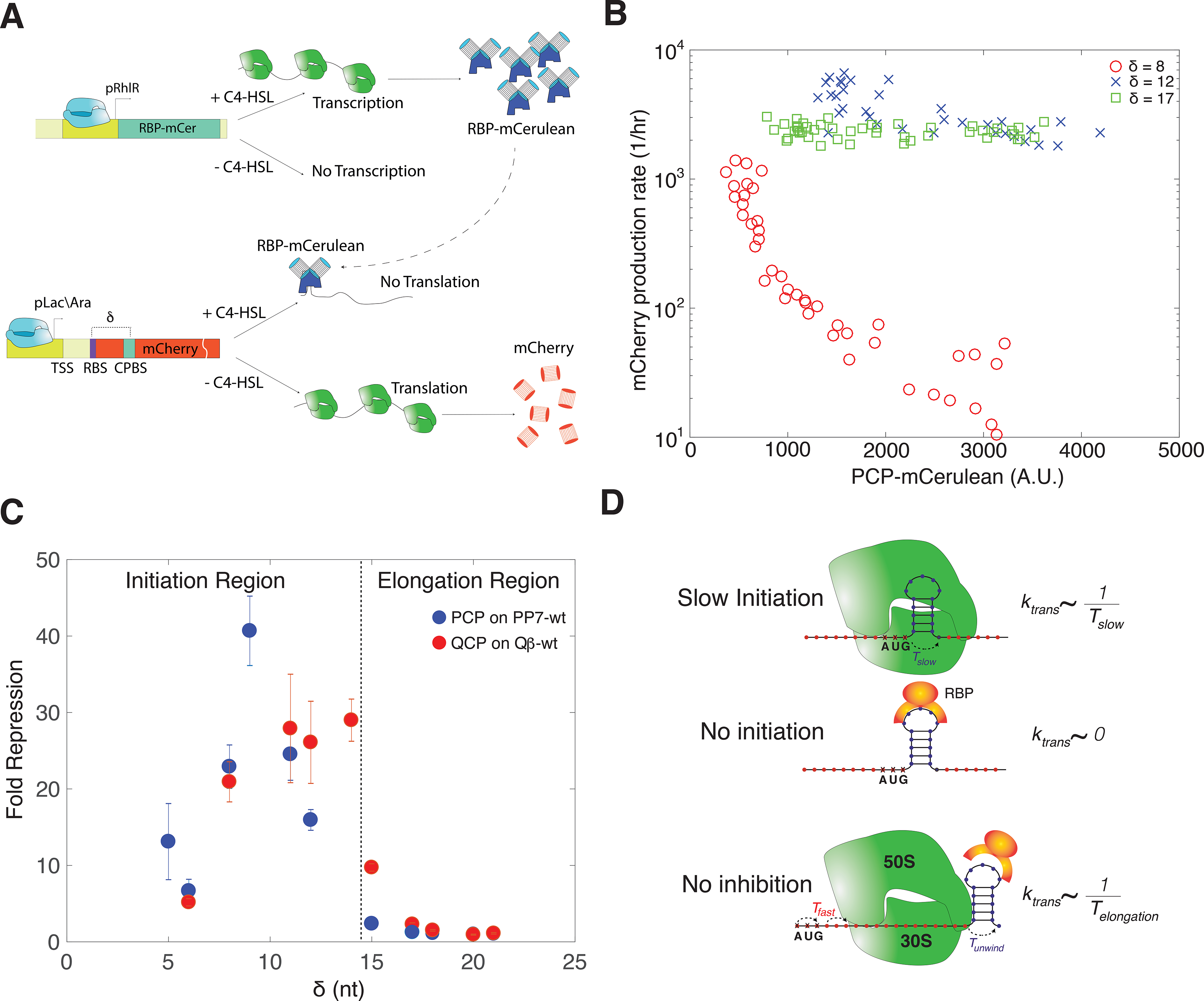
Translational regulation by a RBP-hairpin complex in the ribosomal initiation region. (A) Experimental schematic. Top: plasmid expressing the RBP-CP fusion from a pRhlR inducible promoter. Bottom: a second plasmid expressing the reporter plasmid with the RBP binding site encoded within the 5’ end of the gene (at position δ >0). (B) Dose-response functions for PCP with a reporter mRNA encoding PP7-wt at three positions: δ =8 (red), δ=12 (blue), and δ =17 (green) nt. (C) Fold-repression measurements for PCP (blue) and QCP (red) as a function of hairpin position δ Fold repression is computed by the ratio of the mCherry rate of production at no induction to the rate of production at full induction. Note, for two constructs (QCP with δ =5 and δ =9) the basal levels without induction were too low for fold-repression measurements. (D) A schematic for the mechanistic repression model. If the hairpin is positioned in the initiation region, some positions lead to a slow initiation characterized by *T_slow_*, which in turn leads a partial inhibition of translation. The bound RBP (middle) is able to disrupt initiation, thus inhibiting translation. If the hairpin is positioned downstream of the initiation region (bottom), the 70S subunit is able to assemble and subsequently unwind the RBP-bound structure leading to a cumulative time-scale associated with translation *T_elongation_*.

We hypothesized that the RBP-bound hairpin complex interferes with the initiation process, thus preventing the 70S elongating subunit from forming. To test this model, we measured the mCherry production rate for the PP7-wt and Qβ-wt sites in the presence of PCP and QCP, respectively, for δ varying from 5 to 21 nt. Since the initiation region is thought (33) to extend from the RBS to about 8-10 nucleotides downstream of the AUG (+12 to +14 as in our examples), the chosen range of δ covered both the initiation and elongation regions. Fig. 2C shows that while strong repression is observed for both RBPs within the initiation region, the RBP inhibition effect completely disappears for δ ≥17, for both binding sites. This is consistent with the regulatory effect generated by the hairpins alone (see Fig. 1B-C), which also seems to persist in the initiation region and diminish in the elongation region. Both Fig. 1 and Fig. 2B-C indicate that there may be two forms of inhibition generating the observed repression effect, and both seem to be related to positioning of the hairpin within the translation initiation region. The first (Fig. 2D – top) is a position dependent inhibition that may be related to ribosomal initiation and/or to the rate of hairpin unwinding within the initiation region (see Supplementary Model and Discussion). The second (Fig. 2D-middle) is an initiation region phenomenon, where the bound RBP seems to be inhibiting the formation of the ribosomal complex (i.e. either preventing 30S binding, 70S formation, or some combination thereof). Consequently, initiation of translation seems to be highly susceptible to inhibition via more than one mechanism.

### *In vitro* SHAPE-seq reveal an extended protected region by PCP

To provide a structural perspective on the inhibition mechanism triggered by the RBPs, we employed SHAPE-seq using the acylimidazole reagent 2-methylnicotinic acid imidazolide (NAI), which modifies the 2’ OH of non- or less-structured, accessible RNA nucleotides as found in single-stranded RNA molecules (23). We hypothesized that SHAPE-Seq data can provide a protection foot-print (as in Smola et al. (34)) that is generated when the RBP is bound to its cognate binding site. SHAPE-seq is a next generation sequencing approach (see Materials and Methods and Supplementary Fig. 2 for details), whereby an insight into the structure of an mRNA molecule can be obtained via selective modification of “unprotected” RNA segments. “Unprotected” segments mean single-stranded nucleotides that do not participate in any form of interaction, which include Watson-Crick base-pairing (secondary structure), tertiary interactions (e.g. Hoogsteen base-pairing, G-quadruplex formation, pseudoknots, etc.), and RBP-based interactions. These modifications cause the reverse transcriptase to stall and fall off the RNA strand, leading to a pool of cDNA molecules at varying lengths. Therefore, by counting the number of reads that end in positions along the molecule we can directly measure the number of molecules within this length and can estimate the propensity of this RNA base to be unbound (i.e. single-stranded). To properly quantify such a foot-print, one needs to also account for a background distribution of read lengths that is generated by the propensity of the reverse transcriptase to stall and abort at any nucleotide of a given RNA molecule irrespective of the presence or absence of a modification. Thus, any SHAPE-seq “reaction” must include both a “modified” sample and a “non-modified” control (see Supplementary Fig. 3 and supplementary table 4).

We first applied SHAPE-seq to the PP7-wt δ=6 construct *in vitro* with and without a recombinant PCP protein present in the modification reaction (see Materials and Methods). To decouple the effect of the modification from the underlying reverse transcriptase background for every construct assayed, we employed a signal-to-noise analysis per position on the SHAPE transcript that is normally referred to as “reactivity” (12, 34–39) (see Supplementary Fig. 3 and associated discussion for other definitions of the reactivity (38, 40, 41)). In our version of this observable, we first compute the ratio of the modified to unmodified read counts at each nucleotide, and subsequently subtract one from the result (in similar manner to (41); this and other methods for reactivity calculation have been reviewed by (38)). Values that are bigger than 0 correspond to the propensity of a nucleotide to be modified. Any negative values are set to 0, indicating that the nucleotides at those particular positions do not get modified. For each SHAPE-seq data-set, we checked that the result of the computation yielded data that was not correlated with the unmodified read-count, indicating that the modified and unmodified signals were decoupled. Finally, we used boot-strapping statistics (as in (35, 37)) and Z-factor analysis (as in (34, 39, 42) - see supplementary information) to identify the regions on the RNA molecule where the observed differences between the +RBP and –RBP are statistically significant (equal to or more than three standard deviations).

In Fig. 3A, we present the results for the reactivity analysis carried out on the *in vitro* SHAPE-seq data for the PP7-wt δ=6 construct with (red line) and without (blue line) the presence of a recombinant PCP protein in the reaction solution. Reactivities are presented as a running average over a 10 nt window to eliminate high frequency noise (for further details about the analysis pipeline, see Supplementary Information and Supplementary Fig. 3). The *in vitro* modification experiments were carried out after refolding of the RNA followed by 30 minutes incubation at 37°C with or without the recombinant PCP, and subsequently modified by the SHAPE reagent (i.e. NAI). The plot shows that for the –RBP case (blue line) the reactivity pattern is a varying function of nucleotide position, reflecting a foot-print of some underlying structure. Namely, the segments that are reactive (e.g. −20 to 40 nt range), and those which are not (e.g. 110-140 nt range), indicate non-interacting and highly sequestered nucleotides, respectively.

**Fig. 3:**
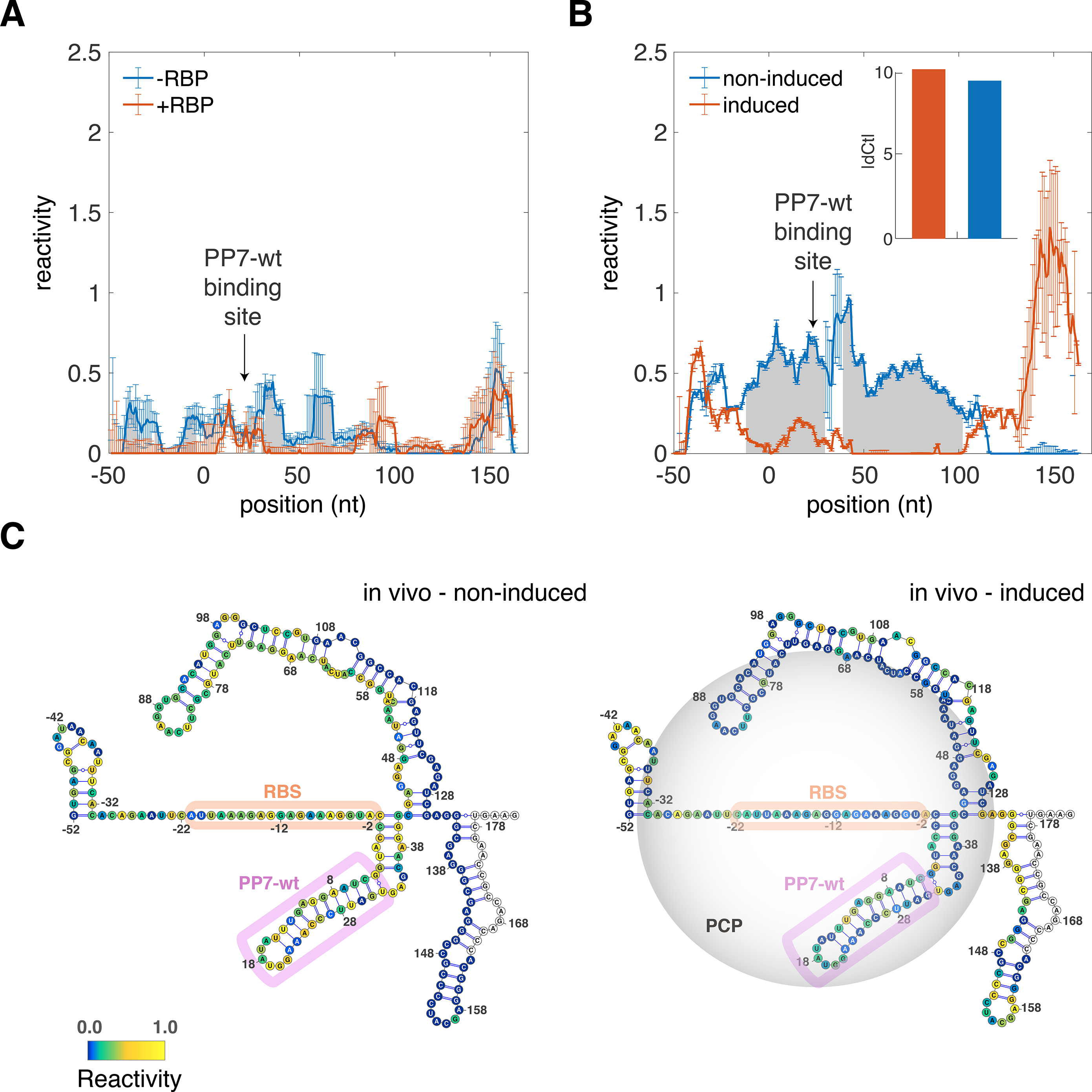
SHAPE-seq analysis of the PP7-wt binding site in the absence and in the presence of RBP (A) *in vitro* reactivity. Scores for the SHAPE-seq reactions carried out on refolded mCherry reporter mRNA molecules containing a PP7-wt binding site at δ=6 with (red) and without (blue) a recombinant PCP present in the reaction buffer. (B) *in vivo* reactivity. Scores for the SHAPE-seq reactions carried out in vivo on the PP7-wt δ=6 construct with the PCP-mCerulean protein non-induced (blue) or induced (red). (Inset) qPCR measurements showing that for both induced and non-induced cases the same level of mRNA was found. For both A and B panels Grey shades signify segments of RNA where a statistically significant difference in reactivity scores (as computed by a Z-factor analysis) was detected between the +RBP and –RBP (A), and induced and non-induced (B) cases respectively. Error-bars were computed using boot-strap resampling and subsequent averaging over two biological replicates. (C) Structural schematics of the segment of the PP7-wt δ=6 construct that was subjected to SHAPE-seq *in vitro*. The structures are overlaid by the reactivity scores (represented as heat-maps from blue, low reactivity, to yellow, high reactivity) for the non-induced (left) and induced (right) cases respectively. Binding site and RBS are highlighted magenta and orange ovals respectively. Grey circle in right structure corresponds to the range of protection by a bound RBP. Non-colored bases correspond to position of the reverse transcriptase primer.

With the addition of the RBP (red line), the reactivity level in the −50 to 80 nt range is predominantly 0 over that range. This indicates that the nucleotides which flank the binding site (positions 6-30 nt) are sequestered and are unmodified or unreactive. We used Z-factor analysis to determine the sequence segments (gray shade) where a statistically significant reduction in reactivity, between the + and – RBP cases, can be observed. These segments span a range ~±50 nts from the position of the binding site, consistent with a previous RNase-based *in vitro* study (43). In contrast, for the positions spanning the range 70-180 nt, the reactivities for both cases are indistinguishable. Together, the reactivity analysis indicates that the RBP is protecting a wide-swath of RNA, which spans the 5’UTR, the initiation, and a portion of the elongation region. This protection is alleviated for positions that are distal from the binding site by > 50 nts, resulting in a realigned reactivity signature indicating that a similar underlying structure for the RNA molecule is maintained for both reaction conditions.

### *In vivo* SHAPE-seq measurements are consistent with *in vitro* measurements

To confirm the observations of the *in vitro* SHAPE-seq protection foot-print, we carried out an *in vivo* SHAPE-seq experiments (see Materials and Methods for differences from the *in vitro* protocol) on the PP7-wt δ=6 construct at two induction states (Fig. 3B): 0 nM of C_4_-HSL (blue line - i.e., no PCP-mCerulean present), and 250 nM of C_4_-HSL (red line – PCP-mCerulean fully induced). The experiments for both conditions were carried in duplicates on different days. To ensure that a proper comparison between the two induction states is carried out, we first checked that the RNA levels at both states were the same using quantitative PCR (Fig. 3Β-inset). We plot in Fig. 3B the reactivity results for both the induced and non-induced cases. For the non-induced case, we observe a strong reactivity signal (>0.5) over the range spanning −45-110 nts, which diminishes to no reactivity for positions > 110. This picture is flipped for the induced case, displaying lower- or no-reactivity for the −40 to 110 nt range and a sharp increase in reactivity for positions > 130 nt. Next, we computed the Z-factor for the regions where the differences between the two reactivity signals was statistically significant (Z>0). In the plot, we marked in grey shades the region where the non-induced reactivity was significantly larger than the induced-reactivity. This shaded region flanks the binding site by ~50 nts both upstream and downstream and is consistent with an interpretation of a wide-swath of PCP protected RNA *in vivo*.

A closer examination of the *in vivo* SHAPE-seq data reveals two major differences from the *in vitro* SHAPE-seq. First, the non-induced case generates significantly higher values of reactivity in the −50-110 nt range as compared with the –RBP *in vitro* case. Second, while in the *in vitro* experiments no significant difference was found between the – and +RBP cases over the 80-180 range, in the *in vivo* case a significant difference was observed. In particular, the non-induced reactivity becomes sharply non-reactive over this range. To gain a structural perspective for the extent of these differences, we plot in Fig. 3C two structures. The structures were computed using RNAfold (44) for the sequence of this molecule and overlaid by its *in vivo* non-induced (left structure) or induced (right structure) reactivity scores (depicted by a heat-map). We demark the RBS (orange oval), PP7-wt binding site (purple oval), and the putative RBP-protected region computed via z-factor analysis (gray circle on right structure). The structures reveal that the reactivity for the non-induced case is inconsistent with the structural prediction. This observation is suggestive of a structure-destabilizing role that an initiating 30S subunit may be generating in the 5’UTR and initiation region. A structural role for the ribosome can be further inferred by the complete lack of reactivity observed in the elongation region of the non-induced case, which is consistent with the presence of a chain of translating ribosomes that may be protecting the RNA from modifications. This is supported by the recovery of the reactivity signal in the elongation region for the induced case, where translation is for the most part abolished. Consequently, the SHAPE-seq analysis *in vivo* reveals significant structural differences between the induced and non-induced cases that are consistent with their RBP-bound states, resultant translational level, and the observed post-transcriptional repression.

### Effective dissociation constant is insensitive to binding-site position

Given the strong RBP-induced repression phenomenon observed in the ribosomal initiation region, we wanted to use this effect to further characterize the binding of the RBPs to structured binding sites. To do so, we first constructed a set of mutated binding sites with various structure-modifying and non-structure-modifying mutations [compare Fig. 4A – bold letters highlighting the native sites for MCP (MS2-wt - top left), PCP (PP7-wt middle-left), and QCP (Qβ-wt bottom-left)]. The mutated binding sites for MCP and PCP were taken from (45, 46) and (9), respectively (Figure 4A), while the ones for QCP were devised by us. All mutations are highlighted in red letters. We then constructed two to four new constructs for each mutated binding site that differed in binding-site position downstream to the AUG. In addition, we constructed a set of control plasmids that lacked a hairpin within the N-terminus of the mCherry reporter gene. Altogether, we constructed 50 reporter plasmids and 5 no-hairpin controls. The new constructs, and the ones previously tested (Fig. 1B and 2B), were co-transformed with all four RBP plasmids to yield 232 RBP—binding-site strains. Our goal with this design was to test not only the binding affinity to the native RBP, but also the relative affinity to the other RBPs, thus obtaining an estimate for the selectivity of RBP binding.

**Fig. 4:**
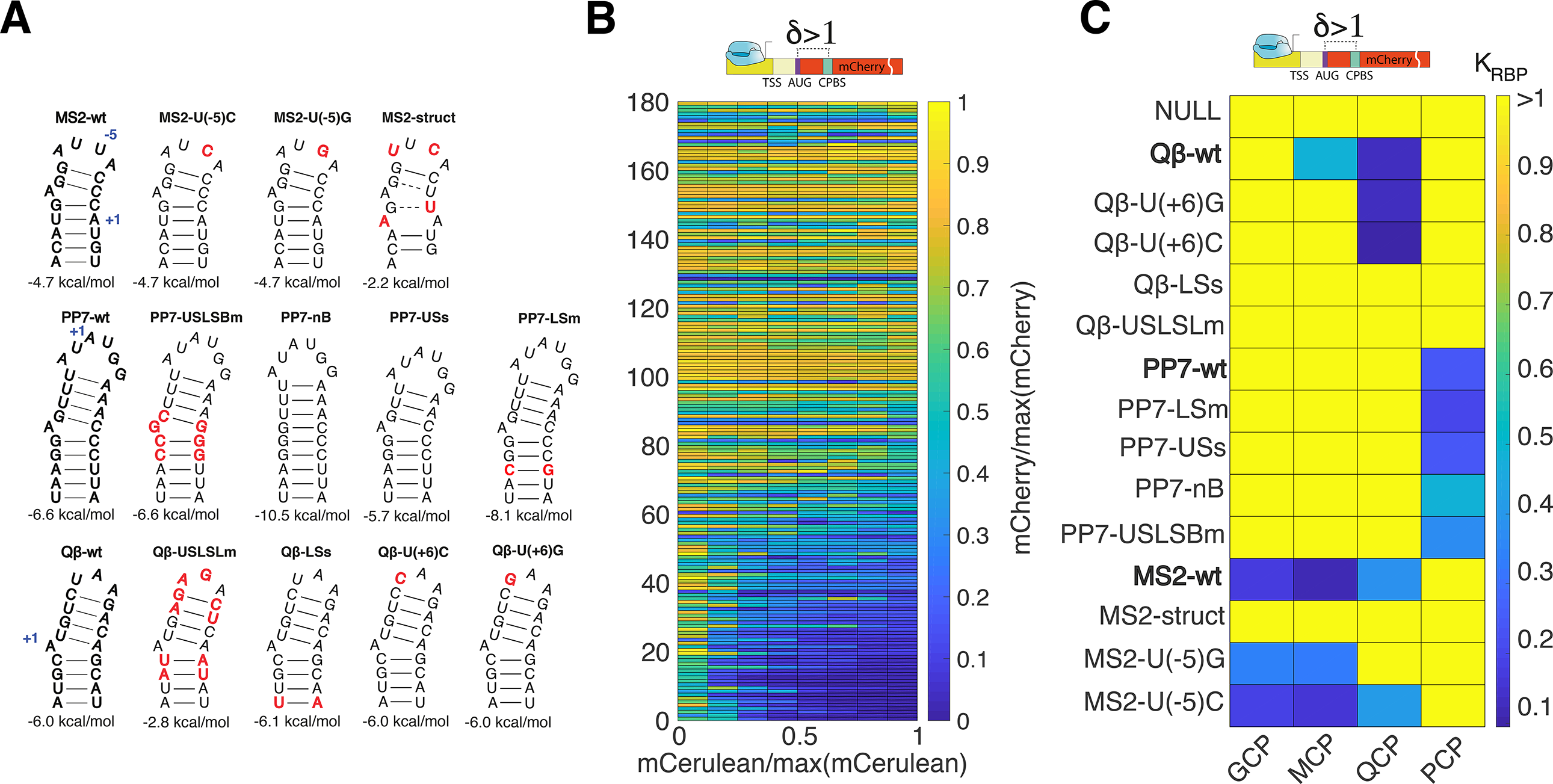
Repression effect can be used to estimate an effective dissociation constant *K_RBP_*. (A)Structural schematic for the 14 binding sites used in the binding affinity study. Red nucleotides indicate mutations from the original wt binding sequence. Abbreviations: US/LS/L/B = upper stem/ lower stem/ loop/ bulge, m = mutation, s = short, struct = significant change in binding site structure. (B) Dose responses for 180 variants whose basal rate of production levels were > 50 a.u./hr. Each response is divided by its maximal mCherry level, for easier comparison. Variants are arranged in order of increasing fold up-regulation. (C) Normalized *K_RBP_* for variants that generated a detectible down-regulatory effect for at least one position. Dark blue corresponds to low *K_RBP_*, while yellow indicates high *KRBP*. If there was no measureable interaction between the RBP and binding site, *K_RBP_* was set to 1.

We plot the dose-response curves of 180 out of the 232 strains as a heat-map in Fig. 4B (strains with basal mCherry rate of production <50 a.u./hr were excluded). In all cases, the data for both the mCherry rate of production and mean mCerulean levels are normalized by the respective maximal value. The dose response functions are arranged in accordance with fold-regulation of the response, with the most repressive variants positioned at the bottom, and the least repressive at the top. The data show that there is a substantial subset of strains, which exhibit strong repression for at least one position (~50 variants), with the strongest mCherry signal occurring at the lowest mCerulean level. To obtain an estimate for the effective binding affinity for each down-regulating variant, we fitted each dose-response curve that exhibited a typical repression response (see Supplementary Figure 1) with a Hill-function-based model (see Supplementary Methods), which assumes a simple relationship between the concentration of RBP measured by its fluorescence, the dissociation constant, and the output expression rate. Finally, we normalized the resulting dissociation constant by the maximal mCerulean expression for the matching RBP to facilitate comparison of the results for the different proteins, yielding an effective dissociation constant (*K_RBP_* – see Supplementary Table 5). Typical error in estimation of the effective dissociation constant was 5-20%, and by averaging *K_RBP_* of each RBP—binding-site pair over multiple positions (values of d) we obtained estimated errors of ~10%.

In Fig. 4C, we plot the averaged *K_RBP_* for different RBP—binding-site combinations as a heat-map, only for those sites (Fig. 4A) for which all four RBPs were tested (“null" corresponds to an average *K_RBP_* computation made on several of the non-binding-site controls). The data show that the effective dissociation constants measured for native sites with their cognate RBPs were low and approximately equal, indicating that native sites are evolutionarily optimized for binding (blue squares). Mutated sites which retained binding affinity displayed slightly larger dissociation constants (light-blue/turquoise), while the *K_RBP_* values of RBP-binding site combinations that did not generate a binding signature were set to the maximum normalized value 1 (*K_RBP_*>1, yellow). When examining the data more closely, we found that PCP is completely orthogonal to the MCP/QCP/GCP group, with no common binding sites. Conversely, we observed crosstalk between the different members of the MCP/QCP/GCP group, with increased overlap between MCP and GCP, which is consistent with previous studies (25).

A closer look at the mutant binding sites reveals that structure-conserving mutations to native binding sites in the loop area [Qβ-U(+6)G, Qβ-U(+6)C, MS2-U(-5)C and MS2-U(-5)G] or stem (PP7-USLSBm and PP7-LSs) did not seem to affect binding of the cognate protein. However, the interaction with a non-cognate RBP is either diminished or eliminated altogether as is the case for MCP with Qβ-U(+6)G and Qβ-U(+6)C, and for QCP with MS2-U(-5)G. In addition, putative structure-altering (MS2-struct – where the lower stem is abolished) and destabilizing (Qb-USLSLm – where the GC base-pairs are converted to UA base-pairs in the lower stem) mutations significantly affected binding. Finally, structure-altering mutations, which retain apparent binding site stability (PP7-nB and PP7-USs), also seemed to retain at least a partial binding affinity to the native RBP. Altogether, these results suggest that binding sites positioned within the initiation region can toler47588888ate multiple mutations as long as certain key structural features necessary for binding and hairpintability (e.g. loop size) are conserved, as was previously observed *in vitro* (9, 46–48).

## DISCUSSION

Synthetic biology approaches have been increasingly used in recent years to map potential regulatory mechanisms of transcriptional and translational regulation, in both eukaryotic and bacterial cells. In this work we built on the work of (22) to quantitatively study RBP-based regulation in bacteria using a combined synthetic biology and SHAPE-seq approach. Using our library of RNA regulatory variants, we were able to identify and characterize a position-dependent repression of translational initiation and elongation by a hairpin structure, and strong repression of initiation when the hairpin was bound by an RBP.

There are multiple layers to our observation of a hairpin regulatory effect within the ribosomal initiation region. First, its very existence has been a source of controversy, with some studies (49–52, 33) suggesting that such structured regions should be detrimental to expression, while others have shown that hairpins do not seem to affect expression in the absence of an RBP (53, 54, 7) or if the stem is shorter than 6 bp. Our structures consist of two stems, 2 to 6 nt long, that are separated by a small 1 nt bulge in the middle. They are characterized by reduced structural stability (e.g. compare PP7-wt and PP7-nb in Figure 4A) that may partially account for the differences from previous observations.

Second, we note the intriguing three nt periodicity in reporter expression finding, which persists throughout the initiation region (i.e. +15 from the AUG) and possibly longer. Since other studies have suggested that ribosomes can unwind hairpin structures (55), it is possible that the observed periodic effect is due to position-dependent resistance to unwinding of the mRNA hairpin during elongation (see supplementary model and Supp. Fig. 4). This explanation is supported by ribosome profiling studies which displayed a trinucleotide periodicity (56, 57), and from several structural studies which found that the first base in a codon is often less structured than the next two bases (40, 58–60). However, given that the effect seems to be localized (or at the very least significantly stronger for δ<15 nt positions) to the initiation region, an additional structural explanation is needed. Previous studies had shown that the 30S subunit can latch on structured mRNA, without having the mRNA bind through the upstream and downstream tunnels (61), and that a small hairpin occupy the A-site without detrimentally affecting elongation (62). Thus, it is possible that this structural scenario is also applicable to our hairpins. In such a case, a trinucleotide periodicity can arise from either a sub-optimal positioning of the hairpin within the A-site, or by varying unwinding rates that are influenced by the position of the hairpin (see model in SI). Consequently, our data indicates that there is still much to be discovered about the regulatory function of hairpins within coding regions, and their relationship to translation initiation and elongation.

When a hairpin exhibiting the 3-nt periodicity signal is located inside the initiation region and its cognate RBP is present at sufficiently high levels, complete repression is observed. However, this repression is sharply attenuated at +10-11 nt from the AUG, and by +16 no RBP-based repression is observed. These results are consistent with the observations of (53, 33), who both showed that structured stems of 6 bp or longer in the N-terminus can silence expression up to +11-13 from the AUG, but show negligible silencing when positioned further downstream. This observation, combined with both the *in vivo* and *in vitro* SHAPE-seq data showing a large swath of protected RNA both upstream and downstream to the binding site, support a model (5, 63–66) in which the bound RBP interrupts initiation (e.g. by obstructing 30S binding, preventing binding of one or more of the initiation factors to the small 30S subunit, inhibiting hairpin unwinding, etc.). However, once the 70S subunit is fully assembled, a hairpin encoded outside the initiation region is unable to inhibit elongation. Interestingly, the *in vivo* SHAPE-seq protection signature measured for PCP spans a larger region of RNA than previously reported both for phage coat proteins *in vitro* (43) and for other proteins with their cognate RNA target using SHAPE-MaP (34). The large protected segment observed here may be a characteristic of phage coat proteins, which suggests a formation of a larger complex that is anchored to the binding site. Regardless, this observation has implications for applications which rely on transcribing cassettes encoding multiple such binding sites and have become common-place in many single molecule in single cell studies published in recent years.

The strong fold repression effect generated by the RBP allowed us to characterize the specific *in vivo* interaction of each RBP—binding-site pair by an effective *K_RBP_*, which we found to be independent of binding site location. Interestingly, the *in vivo K_RBP_* measured for some of the binding sites relative to their native site, differ from past *in vitro* and *in situ* measurements. In particular, PP7-nB, PP7-USs, and MS2-U(-5)G exhibited little or no binding in the *in vitro* setting (9, 46), yet displayed strong binding in our assay; while, MS2-U(-5)C exhibited a reverse behaviour- very high affinity *invitro* and lower affinity in our assay (46). Finally, MS2-struct showed no binding in our assay, but exhibited an affinity higher to that of the wild type in an *in situ* setting (45). These discrepancies may be due to structural constraints, as our *in vivo* RNA constructs were significantly longer than what was used previously *in vitro* and included a 700 nt reporter gene. Another reason for these differences may stem from variations in structure of RNA molecules that stems from their presence inside cells. Our SHAPE-seq analysis revealed that for at least the one construct that was characterized, a translationally active mRNA molecule is less structured *in vivo* as compared with its counterpart *in vitro*. This phenomenon was also previously observed in other studies (67–69). Such structural differences may lead to intramolecular interactions that yield stable folded states *in vivo* that are more amenable to binding as compared with the short constructs that were used in the *in vitro* experiment, and vice versa.

Finally, we found that both MCP and QCP can bind binding sites with different loop sizes than the wild-type binding sites with relatively high affinity. While they do not seem to be sensitive to the sequence content for a loop whose size is equal to the cognate loop (i.e. 4 nt for MCP and 3 nt for QCP), sequence sensitivity is observed for non-cognate loop sizes for both RBPs. This implies that either [GCP, QCP, and PCP] or [MCP, QCP, and PCP], are capable of binding mutually-orthogonal binding sites that differ in structure, opening the door on smart design of mutated binding sites for applications where either set of the three RBPs can be used simultaneously. Our work thus establishes a blueprint for an *in vivo* assay for measuring the dissociation constant of RBPs with respect to their candidate binding sites in a more natural *in vivo* setting. This assay can be used to discover additional binding sites for known RBPs, which could be utilized in synthetic biology applications where multiple non-identical or orthogonal binding sites are needed.

## SUPPLEMENTARY DATA

Supplementary Information file includes detailed modelling and analysis routines.

4 supplementary figures.

5 supplementary tables.

## ACKNOWLEDGEMENT

The authors would like to acknowledge the Technion’s LS&E staff (Tal Katz-Ezov and Anastasia Diviatis) for help with sequencing of the SHAPE-seq fragments.

